# Assessment of the transmural unipolar electrogram morphology change radius during contact force-guided pulmonary vein isolation using the VISITAG™ Module and CARTOREPLAY™

**DOI:** 10.1101/284539

**Authors:** David R. Tomlinson, Kara N. Stevens, Adam J. Streeter

## Abstract

**Aims:** To investigate the radius of transmural (TM) ablation effect at the left atrial posterior wall (LAPW) during contact force (CF)-guided pulmonary vein isolation (PVI), using pure R unipolar electrogram (UE) morphology change – a histologically validated marker of radiofrequency (RF)-induced TM atrial ablation.

**Methods:** Following PVI in 24 consecutive patients (30W, continuous RF), VISITAG™ Module and CARTOREPLAY™ (Biosense Webster Inc.) RF and UE data at left and right-sided LAPW annotated sites 1 and 2 were analysed.

**Results:** Acutely durable PVI without spontaneous / dormant recovery was achieved following 15s and 10-11s RF, at sites 1 and 2, respectively (p<0.0001). At site 1, RS UE morphology was noted pre-ablation, with RF-induced pure R UE morphology change in 47/48 (98%). Left and right-sided second RF site annotation was at 5.8mm and 5.2mm from site 1 respectively (p=0.64), yet immediate pure R UE morphology was noted in 35/48 (73%). For second-annotated sites, 30 demonstrated inter-ablation site transition time ≤17ms; pure R UE morphology was noted at annotation onset in 22/30 (73%), with overall median time to pure R morphology change significantly shorter than at site 1 – 0.0s, versus 4.1s and 5.3s, for left and right-sided first-annotated LAPW sites, respectively (p<0.0001).

**Conclusion:** When the first and second-annotated LAPW RF sites were within 7mm, 73% second-annotated sites demonstrated immediate pure R UE morphology change. These analyses support a paradigm of shorter RF duration at immediately adjacent sites during continuous RF application, and may usefully inform the further development of “tailored” approaches towards CF-guided PVI.

**What’s known?:**

- The VISITAG™ Module and CARTOREPLAY™ permit investigations into the tissue effects of RF energy delivery *in vivo*, via objective annotation methodology and retrospective evaluation of histologically validated unipolar electrogram (UE) criteria for transmural (TM) atrial ablation.
- Greater RF energy effect is seen at left compared to right-sided first-annotated left atrial posterior wall (LAPW) sites during pulmonary vein isolation (PVI).

**What’s new?:**

- Following ∼15s RF delivery at first-annotated LAPW sites and aiming for ≤6mm inter-ablation site distance during continuous RF delivery, 73% second-annotated sites demonstrated immediate TM UE morphology change.
- At second-annotated sites, ∼10s RF resulted in acutely durable PVI in all. Greater left-sided RF energy effect was observed, not explained by differences in RF duration, mean CF or catheter position stability.
- The radius of TM RF effect may be determined at the LAPW following CF and VISITAG™ Module-guided PVI.

## Introduction

The successful completion of electrical pulmonary vein isolation (PVI) for atrial fibrillation (AF) requires a permanent and transmural (TM) ablation lesion encirclement.^1,2^ To achieve this with contact force (CF)-guided radiofrequency (RF) technology, individual lesions must overlap, however excessive local RF delivery may result in life-threatening complications due to extra-cardiac thermal trauma.^3^ Indeed, due to the close proximity of the oesophagus to the left atrial posterior wall (LAPW), inadvertent oesophageal thermal injury may be seen in up to 48% patients.^4^ Despite considerable research efforts to identify suitable RF targets, recent reports of an increase in the rate of atrio-oesophageal fistula following the introduction of CF-sensing catheters^5,6^ indicates a possible causative relationship, and highlights the risks of conducting PVI with an imperfect understanding of when TM ablation has been achieved. Clearly, development of accurate means to assess the occurrence and radius of TM ablative effect is desirable towards achieving “tailored” RF delivery protocols, and should represent an important advance towards improving the safety and efficacy of CF-guided PVI.

Promising recent research has demonstrated greater PVI procedural success using the VISITAG™ Module and ablation index (AI) – an approach with theoretically greater merit in view of the objective RF annotation processes and the incorporation of RF power into the derived ablation targets.^7^ However, this research is not without methodological concerns: (1) During the AI target protocol derivation phase^8^, CF-guided RF ablation was performed employing end-points lacking suitable scientific validation – i.e. bipolar electrogram (BE) abatement of 80%^9^ and RF duration 20-40s.^8,10^ Importantly, a recent report into the utility of near field ultrasound (NFUS) to assess the real-time development of lesion transmurality in canines demonstrated atrial TM RF effect with only 43% BE attenuation (25-30W, 5ml/min), and NFUS illustration of TM effect during LA RF application within 14s.^11^ Furthermore, during PVI employing LAPW ablation at 25W (CF 10-20g), oesophageal temperature alerts >39°C were demonstrated as early as 7s following RF onset^12^; (2) ACCURESP™ adjustment was applied as required to the VISITAG™ Module filters^8^, yet this tool is without *in vivo* validation; (3) The chosen VISITAG™ Module CF filter (force-over-time 30%, minimum 4g) permits intermittent catheter-tissue contact and therefore fails in one important aspect towards fulfilling a suitable definition of positionally stable catheter tip-tissue interaction during RF.^8^ Furthermore, recent *in vivo* data have demonstrated significantly greater RF ablation energy effect at left versus right-sided LAPW sites during PVI; i.e. noted from our earlier work^13^ and also suggested by a report into “forced” ablation power reduction during LAPW ablation according to oesophageal location, where both maximal measured intra-luminal temperature (41°C) and lowest forced power reduction (15.9W) occurred with left-sided oesophageal location compared to central / right-sided course. These data suggest that any single derived target value for LAPW CF-guided RF ablation may incur greater risk of extra-cardiac thermal trauma at left-sided locations, and/or non-TM RF delivery during right-sided ablation.^14^

Previous investigations demonstrated that VISITAG™ Module (Biosense Webster Inc., Diamond Bar, CA) automated RF annotation may be used as an investigational tool following PVI, permitting the development of a short RF duration and VISITAG™-guided CF PVI protocol with excellent efficacy (continuous RF application at 30W, 17ml/min, CF filter force-over-time 100% minimum 1g).^15^ This approach was derived using observations of annotated RF delivery according to the occurrence of intra-procedural recovery of pulmonary vein (PV) conduction; i.e. without reference to electrogram-based tissue response data. However, a subsequent canine study using CF-guided power-controlled irrigated RF at 30W demonstrated the potential utility of real-time pure R unipolar electrogram (UE) morphology change towards the attainment of “tailored” RF; histologically validated TM atrial ablation was achieved in 95% of lesions terminated at the first occurrence of pure R UE, with time to pure R UE, ∼7s and mean lesion depth 4.3mm. Furthermore, while increasing the RF duration to pure R + 5s resulted in histologically TM ablation in 100% without extra-cardiac thermal trauma, lesions terminated at pure R + 10s, +20s and “conventional” 30s total RF, although resulting in 100% pure R UE morphology and histologically proven TM ablation, had 11-17% occurrence of extra-cardiac thermal trauma.^16^

Subsequently, the release of CARTOREPLAY™ (Biosense Webster) permitted assessment of UE morphology change following PVI employing this previously derived CF and VISITAG™-guided PVI protocol. These investigations demonstrated significantly shorter time to pure R UE morphology change at left-sided first-annotated LAPW sites (4.9s v 6.7s; p=0.02), associated with significantly greater impedance drop, compared to right-sided sites. These findings were explained neither by differences in RF duration (∼15s) nor maximum catheter tip distance moved, and occurred despite significantly greater mean contact force (CF) at right-sided sites.^13^ However, when aiming for ≤6mm inter-ablation site distance during continuous RF application, “per protocol” subsequently annotated sites required minimum target RF duration of ∼9s.^15^ Therefore, the purpose of this present report was to retrospectively investigate the onset timing of TM UE morphology change at second-annotated LAPW sites according to the distance from site 1, thereby aiming to identify the radius of TM ablation effect resulting from this previously developed approach to PVI.

## Methods

Retrospective analysis of VISITAG™ Module and CARTOREPLAY™ data was performed at first and second-annotated LAPW ablation sites following single-operator PVI, for the same previously reported consecutive group of unselected patients with symptomatic AF undergoing PVI according to current treatment indications^3^ and as previously descrtibed.^13^

Briefly, all procedures were undertaken using general anaesthesia (GA) and intermittent positive pressure ventilation (IPPV), with temperature-controlled CF-guided RF at 30W delivered using Agilis™ NxT sheath (St Jude Medical Inc., Minneapolis, MA) support. ACCURESP™ Module (Biosense Webster) respiratory training was applied as required to complete the CARTO®3 geometry (V.3, Biosense Webster). VISITAG™ Module filter settings were as previously reported: Positional stability range 2mm, minimum duration 3s; force-over-time 100%, minimum 1g (i.e. constant catheter-tissue contact), without ACCURESP™ Module adjustment. Lesion placement was guided by the VISITAG™ Module, with the preferred site of first RF application at the LAPW opposite each superior PV ∼1cm from the PV ostium; in cases where constant catheter-tissue contact could only be achieved with maximal CF ≥70g, an adjacent LAPW site with lower peak CF was chosen. The target annotated RF duration at each first-annotated LAPW site, as well as any subsequent site ablated from 0W (i.e. “RF ON” sites) and the carina (if ablated) was 15s, whereas ∼9s was the minimum target for all other sites consecutively annotated during continuous RF delivery at 30W; target inter-ablation site distance was ≤6mm. Following completion of circumferential PVI (entrance and exit block), spontaneous recovery of PV conduction was assessed and eliminated during a minimum 20-minute wait; dormant recovery was evaluated and eliminated a minimum of 20 minutes after the last RF. Neither oesophageal luminal temperature monitoring nor post-ablation endoscopic evaluation was employed.

We calculated the catheter tip (position) SD and maximum displacement (from the mean position) at each annotated site from exported VISITAG™ Module data using R 3.3.3.^17^ The RF duration, mean CF, force time integral (FTI) and impedance drop for each annotated site was obtained from exported VISITAG™ Module data; inter-ablation site distance was obtained using the proprietary measurement tool. Retrospective UE analysis was performed as previously described, with the time to pure R UE morphology change taken at the first of 3 consecutively occurring pure R complexes, no more than 1 of which could be an atrial ectopic.^13^ All procedures were performed during proximal pole CS pacing at 600ms. Pictorial evidence of the stated UE morphological changes was provided via a JPEG creation tool (e.g. figures 1A and B); site 1 to 2 transition images for cases 13, 15-16, 19-22 and 24 are shown in the data supplement.

**Figure 1:**
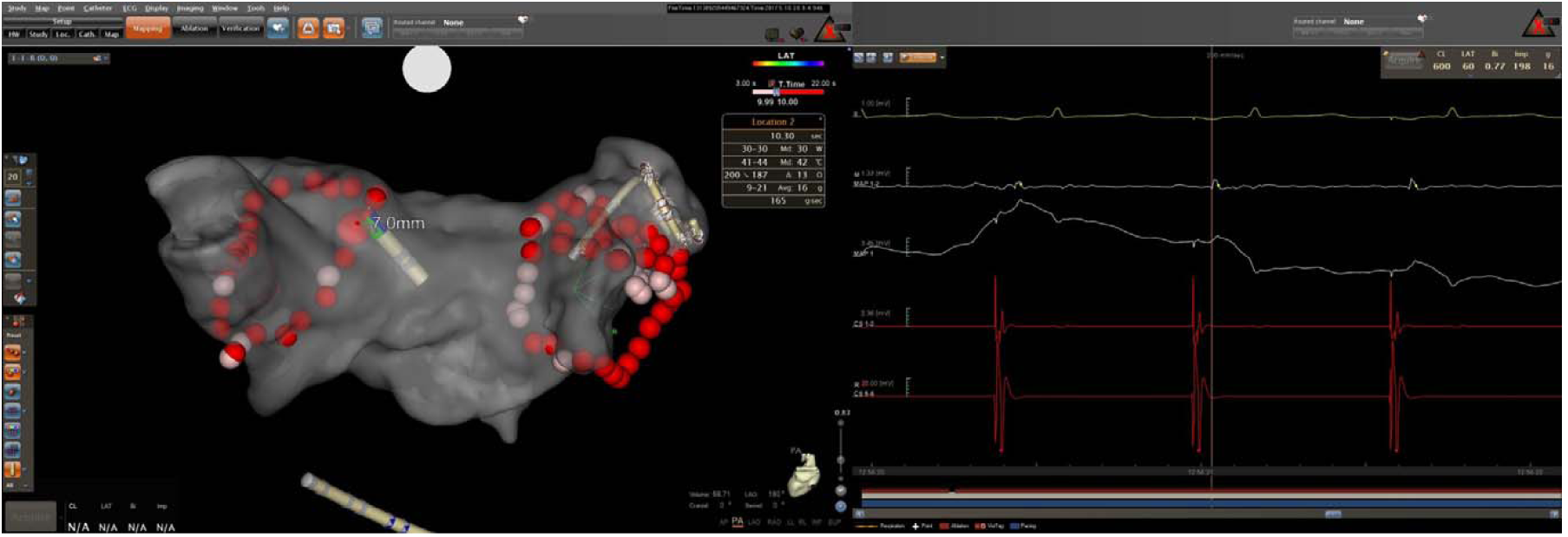
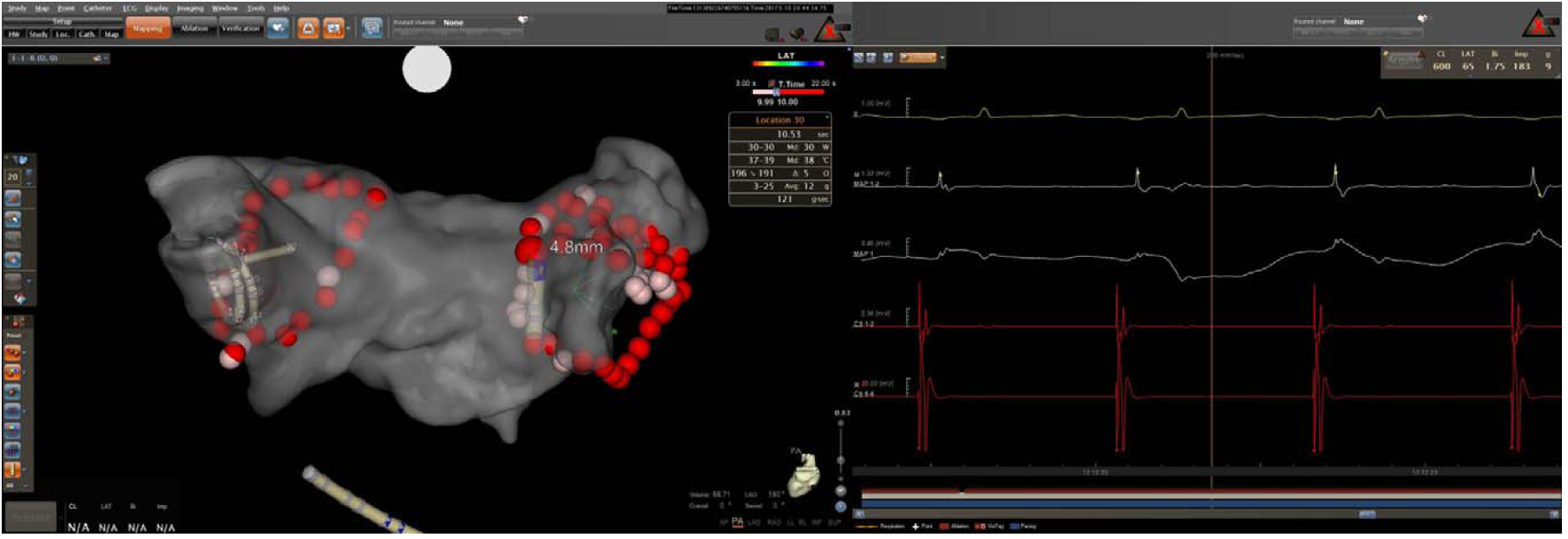
CARTO®3 LA geometry (left panel, transparent PA view) and CARTOREPLAY™ screen (right panel, 200mm/s) at the transition between first and second-annotated sites of RF application in patient #24; site indicated by the gap in the red annotation bar at the bottom of the right panel, indicating 16/17ms interval). VISITAG™ Module ablation site annotation is displayed as 2mm radius spheres, with transition to red at 10s RF duration. (A) Left-sided site 1-2 transition, with “Location 2” highlighted at a distance of 7.0mm from “Location 1” and annotated ablation data shown (box). CARTOREPLAY™ electrogram data demonstrates immediate distal ablation catheter pure R UE morphology during CS pacing at 600ms cycle length (from top to bottom: ECG lead II; MAP 1-2 bipolar electrogram 1.33mV scale; MAP 1 UE 3.45mV scale; CS 1-2 and 5-6 bipolar electrograms); 24-hour clock timeline shown below. (B) Right-sided site 1-2 transition, with “Location 30” highlighted at a distance of 4.8mm from “Location 29” and annotated ablation data shown (box). CARTOREPLAY™ electrogram data demonstrates immediate distal ablation catheter pure R UE morphology during CS pacing at 600ms cycle length.

Statistical analyses were performed using GraphPad Prism version 4.03. D’Agostino and Pearson omnibus normality testing was performed, with parametric data expressed as mean [SD] and non-parametric data as median (1st – 3rd quartile; i.e. inter-quartile range (IQR)). Unpaired / paired t test, Mann Whitney / Wilcoxon signed rank tests were used to assess statistical significance for continuous data, as appropriate. P <0.05 indicated a statistically significant difference. This work received IRB approval for publication as a retrospective service evaluation; all patients provided written, informed consent.

## Results

Twenty-four patients underwent first-time PVI as described, between 24^th^ November 2016 and 11^th^ May 2017. Eighteen were male (75%), with age 58 [SD: 14] years and CHA_2_DS_2_-VASc score 1.4 [SD: 1.3]. AF phenotype was persistent in 13, paroxysmal in 11. Complete PVI was achieved in all without spontaneous / dormant recovery of PV conduction, following 16.1 [SD: 3.1] minutes of RF, without procedural complications.

All 48 first-ablated LAPW annotated sites demonstrated RS UE morphology pre-ablation, with all except 1 left PV site (accidental catheter displacement after 4.4s RF annotation) demonstrating pure R UE morphology change before annotation transition to site 2. At site 2 annotation onset, pure R UE morphology was immediately present in 17/23 (74%) and 18/24 (75%) left and right-sided sites respectively; corresponding inter-ablation site distances were 5.8 (IQR: 5.2-6.2) mm and 5.2 (IQR: 4.9-7.1) mm (p=0.27), and transition times 17 (IQR: 16-1100) ms and 760 (IQR: 17-2258) ms (p=0.002), respectively. At second-annotated sites, pure R UE morphology change was achieved in all left-sided sites, whereas inadvertent catheter displacement at 3.1s resulted in failure to achieve pure R UE morphology at one right-sided site. Overall time to pure R UE morphology change at second-annotated sites was 0.0 (IQR: 0.0-1.2) s and 0.0 (IQR: 0.0-0.3) s for left and right-sided sites, respectively (p=0.90).

To eliminate possible error in the assessment of the timing of pure R UE morphology change onset at second-annotated sites due to “non-annotated” RF delivery (i.e. >2mm positional stability and/or 0g CF events), subsequent analyses excluded all inter-ablation site transitions with >17ms interval (i.e. reflecting the minimum VISITAG™ Module system timing interval). The one instance of failure to achieve pure R UE morphology change at annotated site 1 and one instance at a second-annotated site were also excluded, leaving 30 pairs of first and second-annotated sites (18 left and 12 right-sided); data are shown in table 1.

**Table 1:** Left atrial posterior wall (LAPW) annotated RF data according to left or right-sided location and ablation sequence (i.e. Location 1 / 2, first and second-annotated sites, respectively). Comparison is shown between locations 1 and 2 for each side of the LAPW. Also shown are comparisons between left and right-sided locations 1 (Loc 1) and 2 (Loc 2). Data is shown as mean [SD] / median (IQR), as appropriate; RF, radiofrequency; CF, contact force; FTI, force time integral; SD, standard deviation; UE, unipolar electrogram.

|  | Left-sided sites |  |  | Right-sided sites |  |  | Left v right |  |
| --- | --- | --- | --- | --- | --- | --- | --- | --- |
|  | *Location 1* | *Location 2* | *p* | *Location 1* | *Location 2* | *p* | *Loc 1: p* | *Loc 2: p* |
| RF duration (s) | 14.9<br>(14.3-15.7) | 9.6<br>(6.1-11.4) | <0.0001 | 15.1<br>(14.8-15.7) | 10.6<br>(10.4-11.3) | <0.0001 | 0.37 | 0.22 |
| Mean CF (g) | 11.7<br>(9.2-13.4) | 14.5<br>(11.7-16.5) | 0.058 | 16.2<br>(12.2-17.7) | 14.2<br>(11.8-19.0) | 0.93 | 0.06 | 0.79 |
| FTI (gs) | 170<br>(127-221) | 134<br>(74-150) | 0.006 | 244<br>(184-270) | 158<br>(115-197) | 0.03 | 0.07 | 0.13 |
| Impedance drop ($\Omega$) | 16.9<br>[8.6] | 7.2<br>[3.7] | 0.0004 | 10.4<br>[2.2] | 4.4<br>[1.5] | <0.0001 | 0.02 | 0.03 |
| Catheter position SD (mm) | 1.4<br>[0.3] | 1.8<br>[0.3] | <0.0001 | 0.9<br>[0.3] | 1.4<br>[0.4] | 0.004 | 0.001 | 0.001 |
| Maximum distance moved (mm) | 2.7<br>[0.5] | 4.4<br>[1.0] | <0.0001 | 2.7<br>[0.7] | 3.4<br>[0.7] | 0.01 | 0.88 | 0.007 |
| Time to pure R UE (s) | 4.1<br>(3.3-6.8) | 0.0<br>(0.0-1.4) | <0.0001 | 5.3<br>(4.4-5.8) | 0.0<br>(0.0-0.7) | <0.0001 | 0.20 | 0.83 |

RF duration at first-annotated LAPW sites was significantly greater than at second-annotated sites, as per the ablation protocol. CF was without significant difference, but FTI was significantly greater at site 1 (i.e. secondary to the greater protocol RF duration). Similarly, impedance drop was significantly greater at site 1, but importantly there was a significantly greater impedance drop at each left-sided site, explained neither on the basis of differences in RF duration, CF, nor measures of ablation catheter positional stability. Indeed, for both catheter position SD and maximum distance moved, left-sided annotated sites demonstrated significantly greater measures of catheter position instability. Notably, the time to pure R UE morphology change was significantly shorter at second-annotated sites (p<0.001). Furthermore, pure R UE morphology was present at second-annotated site onset in 13 of 18 (72%) left-sided and 9 of 12 (75%) right-sided sites (figure 2); overall 22/30 (73%). The single outlying second site with time to pure R UE morphology 6.9s (at 4.1mm distance from site 1), had corresponding first site RF duration 15.2s, mean CF 7.7g, FTI 117gs, impedance drop 26Ω and time to pure R UE morphology 9.5s.

**Figure 2:**
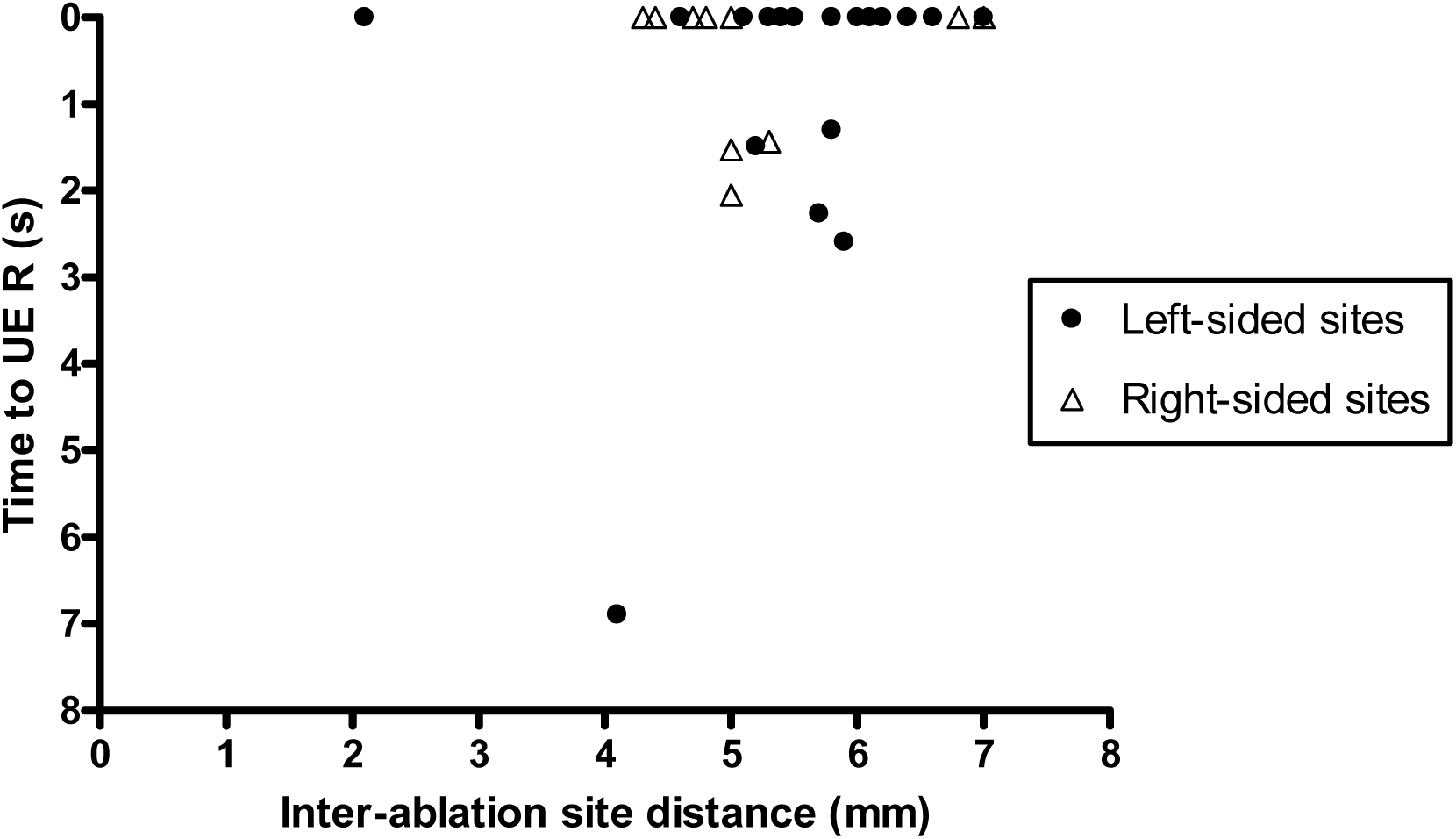
Time to pure R UE morphology change at second-annotated sites (“UE R”) demonstrating ≤17ms inter-ablation site transition, according to the distance from annotated site 1; UE R scale is reversed to better visualise data indicating 0s time to achieve pure R UE morphology change.

## Discussion

This study presents a description of novel methodology permitting the *in vivo* assessment of the radius of RF-induced thermal effects during PVI, utilising VISITAG™ Module objective ablation site annotation and CARTOREPLAY™. Pure R UE morphology change – a histologically validated marker of TM atrial ablation^16,18^ – was analysed at first and second-annotated sites of RF delivery during LAPW ablation employing continuous RF delivery. Together with measured inter-ablation site distances, these data identified a radius of thermal injury up to 7mm from annotated site 1 (figure 2); 73% second-annotated sites demonstrated immediate TM UE morphology change, with the time to pure R UE morphology change significantly shorter overall. Furthermore, the demonstration of acutely durable PVI resulting from significantly shorter “per protocol” RF delivery at second-annotated sites, indicates that when PVI is performed in the manner described, continuous RF delivery is likely to represent a more efficient approach towards TM linear lesion creation than point-by-point (PbP) RF application. This hypothesis is supported by findings from an *ex vivo* bovine myocardial model of linear ablation, where compared to PbP ablation, significantly greater lesion volumes resulted from continuous RF application despite equivalent energy delivery. These findings were explained on the basis of continuous RF not permitting the passive convective cooling that may be apparent between interrupted RF applications.^19^ A subsequent *in vivo* magnetic resonance imaging (MRI) thermography study undertaken during canine LV RF application demonstrated an expanding thermal lesion as power was increased.^20^ Extrapolating these data to the left atrium during PVI, it may be postulated that the small catheter tip movements during VISITAG™ Module guidance in our study resulted in new ablation site annotation, yet remained within a zone of thermal injury secondary to RF application at site 1. Theoretically, durable RF effect at such second-annotated sites could be achieved using significantly shorter RF delivery considering the higher starting tissue temperature, evidenced by the immediately apparent UE morphology change. This finding is clearly of great clinical relevance in view of the common practice of continuous RF delivery during RF, yet with presently no adjustment towards RF delivery at consecutive sites suggested in CF-guided PVI “guideline recommendations”.^21,22^ Indeed, an awareness of these novel findings may facilitate a reduction in RF delivery without compromising procedural efficacy, while potentially helping to reduce the risk of extra-cardiac thermal trauma.

This present report also confirms the previous finding of greater RF energy effect at left-sided LAPW sites *in vivo*, and extends this to second-annotated sites during continuous RF application. This cannot be explained by differences in measured RF delivery (RF duration, CF, power), and perhaps counter-intuitively was noted despite significantly greater measured ablation catheter positional instability at left-sided sites. Compared to first-annotated sites, there was also greater positional instability at all second-annotated sites; however, this can be explained by the fact that second-annotated sites by definition included one deliberate catheter tip acceleration / deceleration event (i.e. at “annotation start”) and another acceleration event (i.e. at “annotation end”), whereas at site 1 there was only one deliberate catheter acceleration event at “annotation end”. The mechanisms underlying greater left-sided LAPW RF energy effect remain unknown, but logically must represent either significantly greater surface area of ablation catheter tip-tissue interaction (and consequent greater energy delivery^23^), and/or less out-of-phase catheter-tissue motion, during left-sided LAPW RF delivery.

Finally, these present data represent electrogram-based evidence supporting the previously employed methodology resulting in the development of a CF and VISITAG™ Module-guided approach to PVI, using continuous RF application.^15^ When combined with the methods for UE morphology assessment described in this present report and taken alongside our suggestion that through the adoption of suitable stability filters, the VISITAG™ Module provides a “language” with which to describe the RF “effector arm”^13^, this approach represents a new methodological platform for *in vivo* RF ablation research in human subjects. Furthermore, given the theoretical universal applicability of such VISITAG™ Module-based analyses, these methods could also be employed during investigations using animal models of CF-guided RF ablation; e.g. investigating the acute tissue response following very short duration, high power delivery. Using this approach and when combined with histologically validated TM tissue RF effects, VISITAG™ Module-based measures of RF delivery and ablation catheter stability of greater scientific validity could be identified and applied in future clinical studies of human subjects.

### Study limitations

This present report is of a single-operator’s practice, utilising GA (with IPPV) in all. Therefore, the described approach to ablation is not a direct recommendation for others to follow without careful consideration. Indeed, in view of the very short second-annotated site time to pure R UE morphology change demonstrated, yet “per protocol” annotated site 2 RF duration of ∼9s, some LAPW lesions were probably accompanied by inadvertent extra-cardiac thermal trauma. Accordingly, it may be possible to achieve durable PVI at lower risk of extra-cardiac thermal trauma through employing even shorter RF duration (and possibly also lower RF power) than reported here; this clearly requires further study. Also, this report is limited to the first two annotated LAPW sites – chosen as this represented the simplest methodology to permit a description of RF-induced changes at immediately adjacent, ablation naive sites. A description of UE morphology change at all VISITAG™ Module-annotated sites during CF-guided PVI would be extremely useful, but is a more complex undertaking and was beyond the scope of this present report. However, these data have already been collected for this same patient cohort and will form the basis of a future report.

Previous single-centre research conducted before the advent of CF sensing demonstrated the potentially reversible nature of pure R UE morphology change at sites of intra-procedural recovery of PV conduction.^24^ Findings from a study of superfused guinea pig papillary muscle responses to thermal insult may provide insights into this finding; i.e. temperature-dependent depolarisation displayed a sigmoidal response, with loss of excitability at >42.7°C, but with reversibility of these findings at temperatures <51.3°C.^25^ Furthermore, our own previous investigations have demonstrated intra-procedural recovery of PV conduction when an inter-ablation site distance ≥6mm was associated with adjacent annotated RF duration of 3.0s.^15^ Although representing an important caveat towards the interpretation of pure R UE morphology change *in vivo*, particularly when this present report was without histological examination of sites of RF delivery, there were no instances of intra-procedural recovery of conduction following RF protocol completion. Furthermore, in view of the second-annotated site “per protocol” RF duration of ∼9s, yet with near instantaneous pure R UE morphology change evident, most second-site RF applications were to at least that duration employed by Bortone *et al* when achieving single-site 100% histologically-confirmed TM lesions of 4.3mm depth.^16^ Although it is scientifically invalid to draw firm conclusions from such comparisons (particularly when VISITAG™ Module annotation was not employed during these animal studies), together with the previously reported high procedural efficacy of this present CF and VISITAG™ Module-guided protocol^15^, it is very unlikely that RF application at these LAPW sites resulted in non-TM lesions.

All UE morphology data were collected retrospectively, so this report does not prove that modification of RF delivery based on real-time UE morphological assessment during VISITAG™ Module and CF-guided PVI is appropriate. Indeed, taken together with the finding of pure R UE morphology change reversibility during non-CF guided PVI^24^, reliance on this factor alone is probably inappropriate. Instead, a more suitable approach may utilise real-time UE morphology change and other measures of energy delivery and/or tissue response (i.e. RF duration, impedance drop) at objectively annotated sites. Obviously, this remains to be proven but is worthy of further research, not least since this may result in the development of “tailored” and possibly safer, RF PVI ablation protocols.

When considering the methodology for timing of pure R UE morphology change onset utilised in this present report, there was possible error introduced according to the chosen base atrial pacing rate. For example and taking the worst possible scenario, the first paced atrial electrogram demonstrating pure R morphology change may have occurred 600ms after second-annotated site onset, yet for simplicity and to avoid far more time-consuming measurements, for reporting purposes this was classified as 0s. However, this should not seriously undermine the overall findings of this present report, since it is highly unlikely that all these intervals were 600ms. Furthermore, even if they were, re-classifying all “0s events” to 600ms would still represent a significantly shorter time to pure R UE morphology change at second-annotated sites.

Finally, the ability of VISITAG™ Module-based RF annotation to function as a “language” for the RF “effector arm” *in vivo* for both research and clinical practice, as suggested, remains to be proven. However, this is an important area for future investigations since without such facility there is no suitable means to standardise and report approaches to RF delivery during future randomised controlled trials, making the task of translating any useful findings into clinical practice, impossible.

## Conclusions

Following CF and VISITAG™ Module-guided RF delivery at first-annotated LAPW sites employing continuous RF delivery, immediate occurrence of pure R UE morphology change at 73% of second-annotated sites indicated ∼7mm radius of thermal injury. Furthermore, acutely durable PVI was achieved with shorter RF duration at second-annotated sites, indicating a theoretical advantage of continuous RF application during linear lesion creation *in vivo*. These novel investigational methodologies may facilitate the development of more efficient and potentially safer “tailored” CF and VISITAG™ Module-guided PVI protocols.

## Acknowledgments

I am grateful to Cherith Wood, Daniel Newcomb and Ian Lines, Cardiac Physiologists, for their technical support into all cases conducted during this report. I am also grateful to Robert Pearce and Vicky Healey (Biosense Webster) for additional technical assistance and to Noam Seker-Gafni, Tal Bar-on, Einav Geffen, Assaf Rubissa and colleagues at the Haifa Technology Center, Israel (Biosense Webster) for their original help with VISITAG™ Module technical queries.

