## Supplementary Materials for "Assessment of the transmural unipolar electrogram morphology change radius during contact force-guided pulmonary vein isolation using the VISITAG™ Module and CARTOREPLAY™"

**Supplementary data**

**Figures demonstrating CARTOREPLAY™ unipolar electrogram morphology data and CARTO^®^3 geometry at the transition between first and second VISITAG™ Module annotated sites**

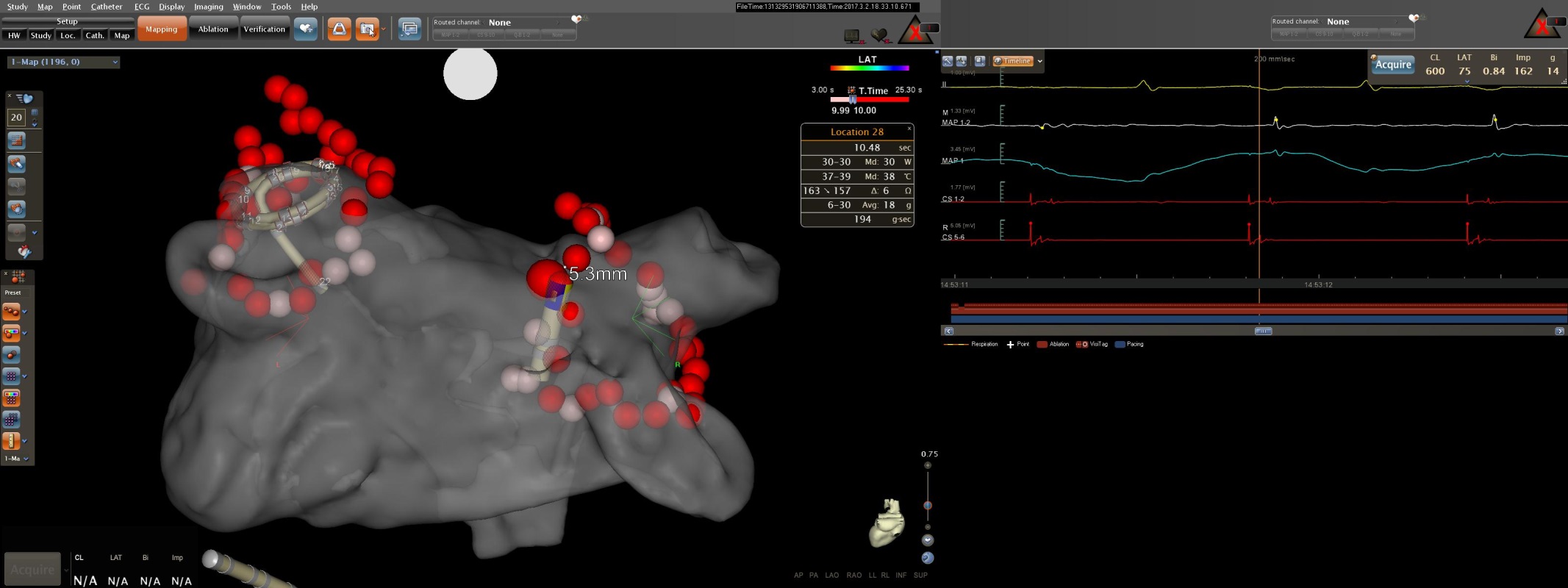

Figure 1: CARTO^®^3 LA geometry (left panel, transparent PA view) and CARTOREPLAY™ screen (right panel) at the transition to the second annotated RF application during right PV isolation in patient #13 (interval = 17ms indicated by the gap in the red annotation bar at the bottom of the right panel). VISITAG™ Module ablation site annotation is displayed as 2mm radius spheres, with transition to red at 10s RF duration; “Location 28” is highlighted with annotated ablation data shown (box), at a distance of 5.3mm from “Location 27”. CARTOREPLAY™ electrogram data at 200ms sweep speed demonstrates RS UE morphology of the second distal ablation catheter electrogram during CS pacing at 600ms cycle length, with transition to the first of 3 consecutive pure R morphology complexes at 1.44s following annotation onset (from top to bottom: ECG lead II; MAP 1-2 bipolar electrogram 1.33mV scale; MAP 1 UE 3.45mV scale; CS 1-2 and 5-6 bipolar electrograms); 24-hour clock timeline shown below.

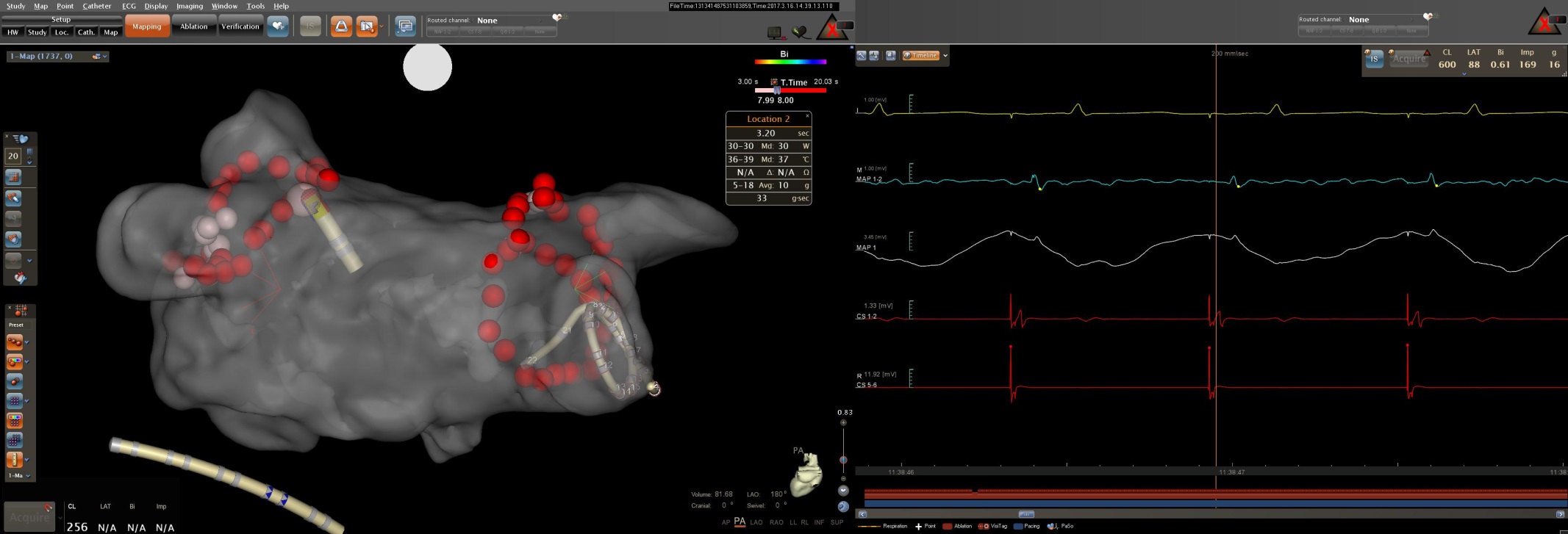

Figure 2A: CARTO^®^3 LA geometry (left panel, transparent PA view) and CARTOREPLAY™ screen (right panel) at the transition to the second annotated RF application during left PV isolation in patient #15 (interval = 17ms indicated by the gap in the red annotation bar at the bottom of the right panel). VISITAG™ Module ablation site annotation is displayed as 2mm radius spheres, with transition to red at 10s RF duration; “Location 2” is highlighted with annotated ablation data shown (box), at a distance of 6.1mm from “Location 1”. CARTOREPLAY™ electrogram data at 200ms sweep speed demonstrates immediate pure R UE morphology of the distal ablation catheter electrogram during CS pacing at 600ms cycle length (from top to bottom: ECG lead II; MAP 1-2 bipolar electrogram 1.00mV scale; MAP 1 UE 3.45mV scale; CS 1-2 and 5-6 bipolar electrograms); 24-hour clock timeline shown below.

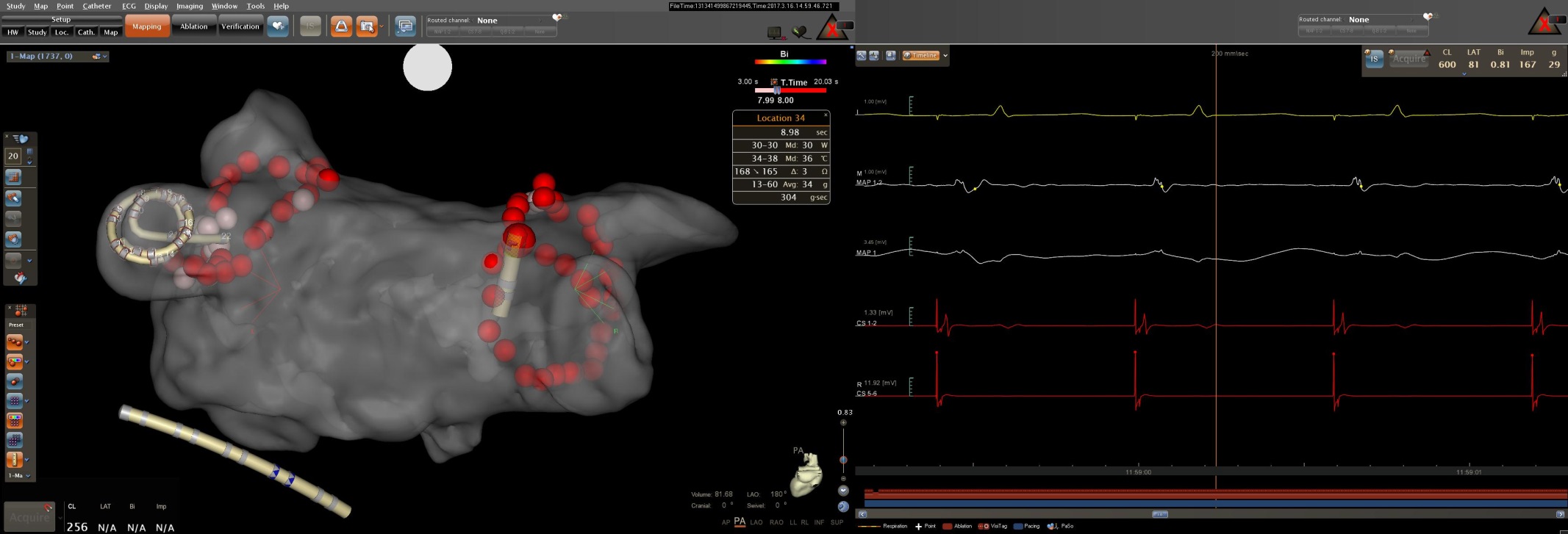

Figure 2B: CARTO^®^3 LA geometry (left panel, transparent PA view) and CARTOREPLAY™ screen (right panel) at the transition to the second annotated RF application during right PV isolation in patient #15 (interval = 17ms indicated by the gap in the red annotation bar at the bottom of the right panel). VISITAG™ Module ablation site annotation is displayed as 2mm radius spheres, with transition to red at 10s RF duration; “Location 34” is highlighted with annotated ablation data shown (box), at a distance of 6.8mm from “Location 33”. CARTOREPLAY™ electrogram data at 200ms sweep speed demonstrates immediate pure R UE morphology of the distal ablation catheter electrogram during CS pacing at 600ms cycle length (from top to bottom: ECG lead II; MAP 1-2 bipolar electrogram 1.00mV scale; MAP 1 UE 3.45mV scale; CS 1-2 and 5-6 bipolar electrograms); 24-hour clock timeline shown below.

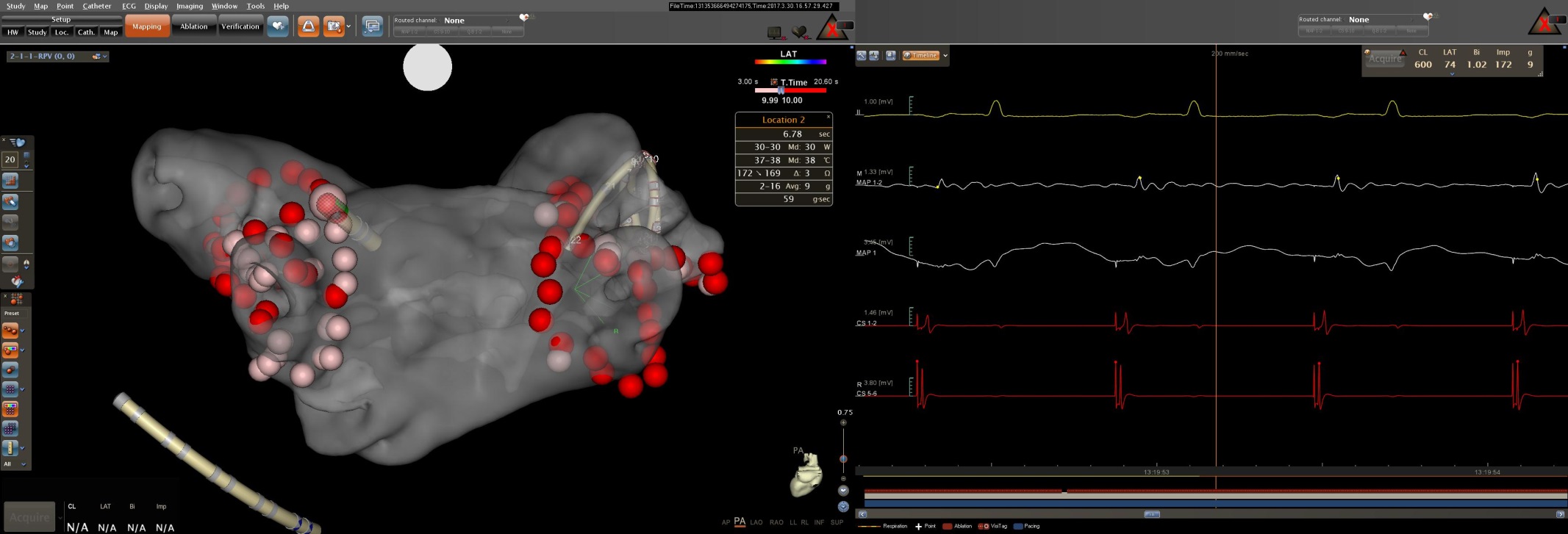

Figure 3A: CARTO^®^3 LA geometry (left panel, transparent PA view) and CARTOREPLAY™ screen (right panel) at the transition to the second annotated RF application during left PV isolation in patient #16 (interval = 16ms indicated by the gap in the red annotation bar at the bottom of the right panel). VISITAG™ Module ablation site annotation is displayed as 2mm radius spheres, with transition to red at 10s RF duration; “Location 2” is highlighted with annotated ablation data shown (box), at a distance of 2.1mm from “Location 1”. CARTOREPLAY™ electrogram data at 200ms sweep speed demonstrates immediate pure R UE morphology of the distal ablation catheter electrogram during CS pacing at 600ms cycle length (from top to bottom: ECG lead II; MAP 1-2 bipolar electrogram 1.33mV scale; MAP 1 UE 3.45mV scale; CS 1-2 and 5-6 bipolar electrograms); 24-hour clock timeline shown below.

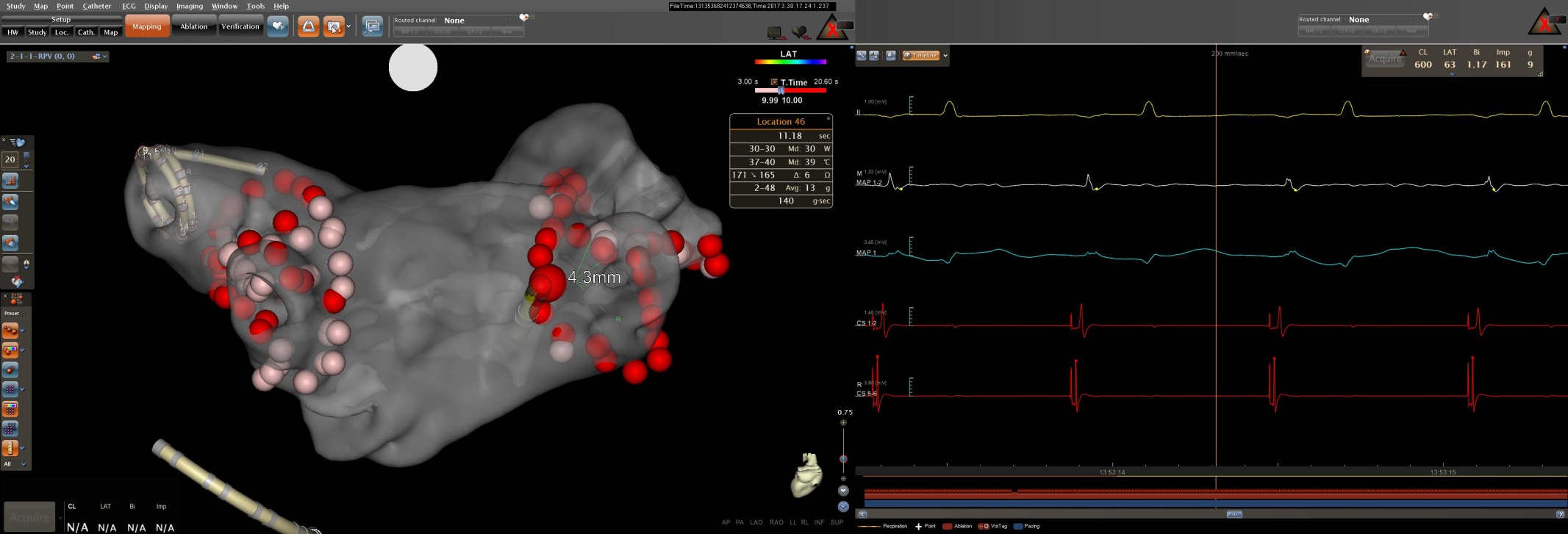

Figure 3B: CARTO^®^3 LA geometry (left panel, transparent PA view) and CARTOREPLAY™ screen (right panel) at the transition to the second annotated RF application during right PV isolation in patient #15 (interval = 17ms indicated by the gap in the red annotation bar at the bottom of the right panel). VISITAG™ Module ablation site annotation is displayed as 2mm radius spheres, with transition to red at 10s RF duration; “Location 46” is highlighted with annotated ablation data shown (box), at a distance of 4.3mm from “Location 45”. CARTOREPLAY™ electrogram data at 200ms sweep speed demonstrates immediate pure R UE morphology of the distal ablation catheter electrogram during CS pacing at 600ms cycle length (from top to bottom: ECG lead II; MAP 1-2 bipolar electrogram 1.33mV scale; MAP 1 UE 3.45mV scale; CS 1-2 and 5-6 bipolar electrograms); 24-hour clock timeline shown below.

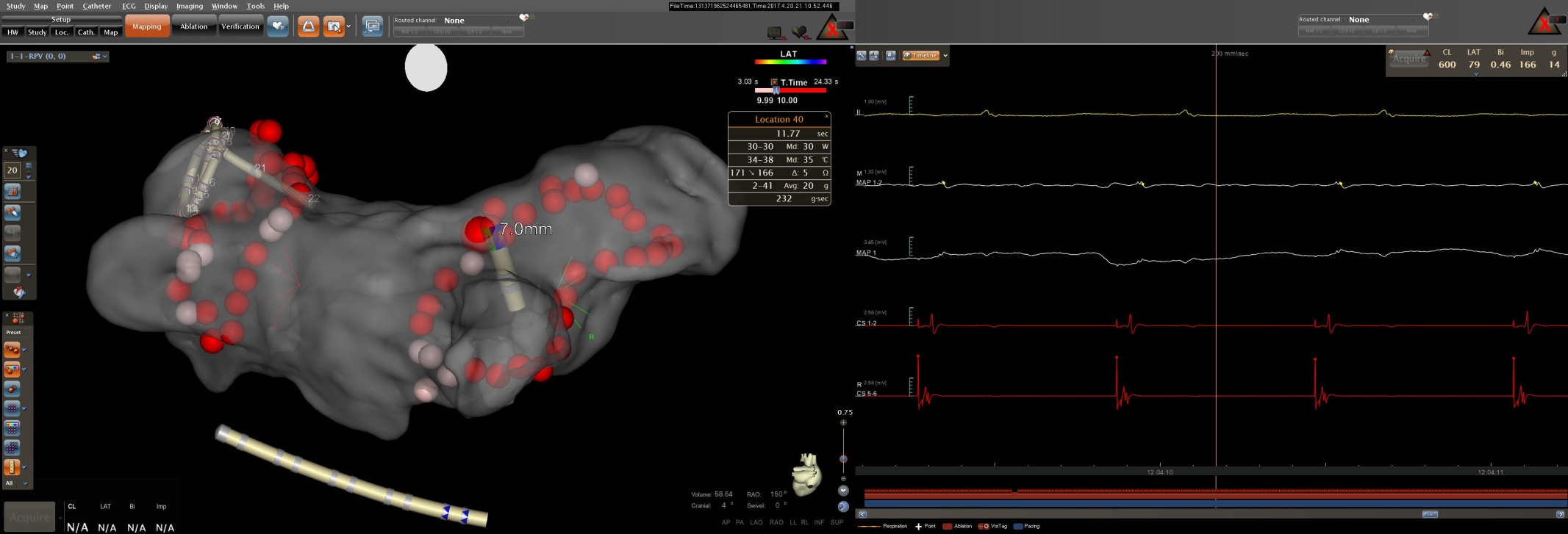

Figure 4: CARTO^®^3 LA geometry (left panel, transparent PA view) and CARTOREPLAY™ screen (right panel) at the transition to the second annotated RF application during right PV isolation in patient #19 (interval = 17ms indicated by the gap in the red annotation bar at the bottom of the right panel). VISITAG™ Module ablation site annotation is displayed as 2mm radius spheres, with transition to red at 10s RF duration; “Location 40” is highlighted with annotated ablation data shown (box), at a distance of 7.0mm from “Location 39”. CARTOREPLAY™ electrogram data at 200ms sweep speed demonstrates immediate pure R UE morphology of the distal ablation catheter electrogram during CS pacing at 600ms cycle length (from top to bottom: ECG lead II; MAP 1-2 bipolar electrogram 1.33mV scale; MAP 1 UE 3.45mV scale; CS 1-2 and 5-6 bipolar electrograms); 24-hour clock timeline shown below.

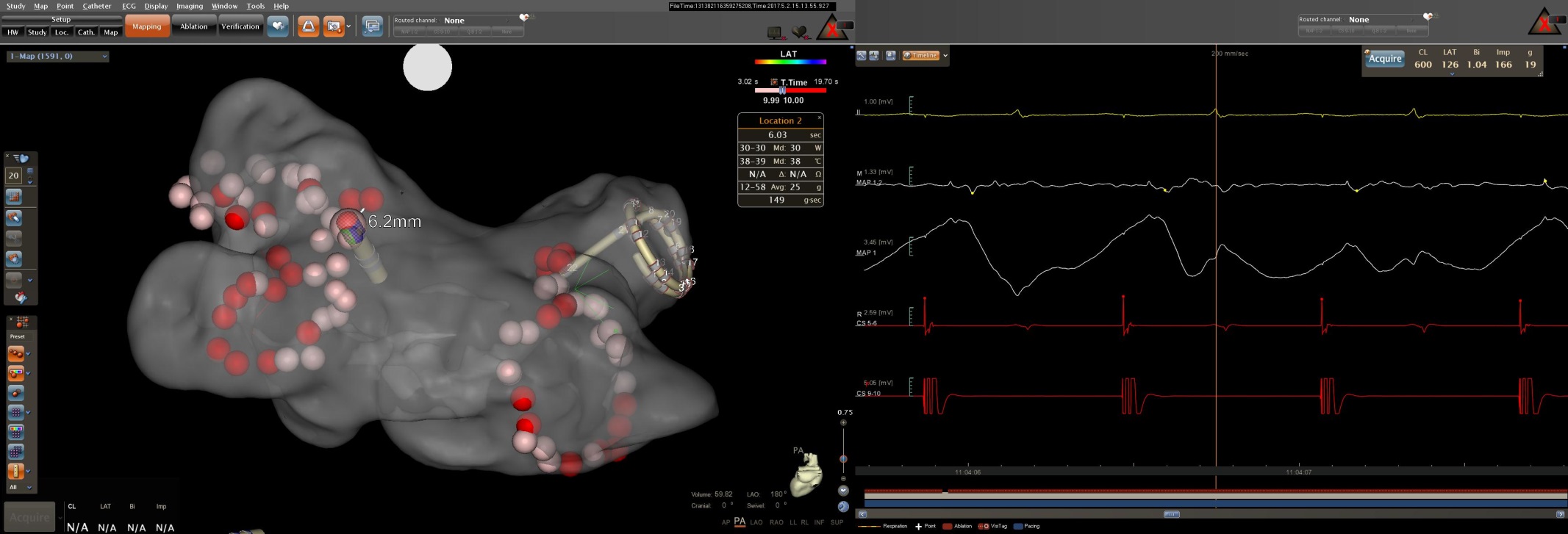

Figure 5: CARTO^®^3 LA geometry (left panel, transparent PA view) and CARTOREPLAY™ screen (right panel) at the transition to the second annotated RF application during left PV isolation in patient #20 (interval = 17ms indicated by the gap in the red annotation bar at the bottom of the right panel). VISITAG™ Module ablation site annotation is displayed as 2mm radius spheres, with transition to red at 10s RF duration; “Location 2” is highlighted with annotated ablation data shown (box), at a distance of 6.2mm from “Location 1”. CARTOREPLAY™ electrogram data at 200ms sweep speed demonstrates some artefact, but with immediate pure R UE morphology of the distal ablation catheter electrogram during CS pacing at 600ms cycle length still clearly visible (from top to bottom: ECG lead II; MAP 1-2 bipolar electrogram 1.33mV scale; MAP 1 UE 3.45mV scale; CS 1-2 and 5-6 bipolar electrograms); 24-hour clock timeline shown below.

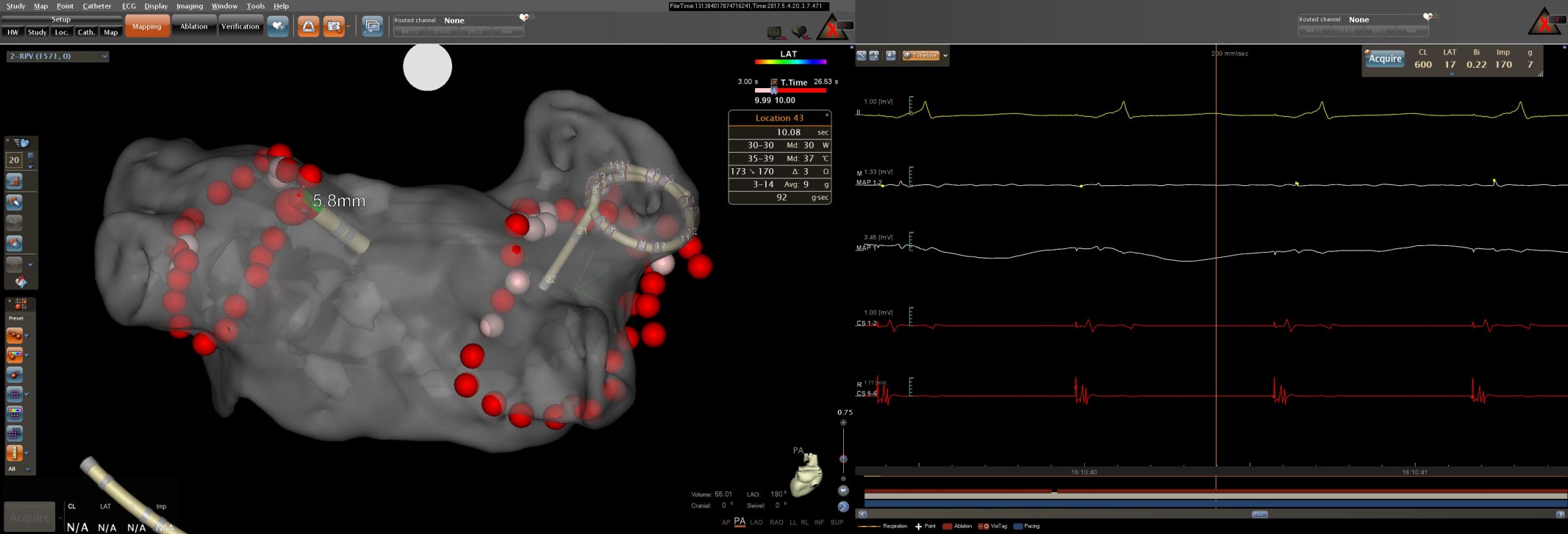

Figure 6A: CARTO^®^3 LA geometry (left panel, transparent PA view) and CARTOREPLAY™ screen (right panel) at the transition to the second annotated RF application during left PV isolation in patient #21 (interval = 17ms indicated by the gap in the red annotation bar at the bottom of the right panel). VISITAG™ Module ablation site annotation is displayed as 2mm radius spheres, with transition to red at 10s RF duration; “Location 2” is highlighted with annotated ablation data shown (box), at a distance of 5.8mm from “Location 1”. CARTOREPLAY™ electrogram data at 200ms sweep speed demonstrates immediate pure R UE morphology of the distal ablation catheter electrogram during CS pacing at 600ms cycle length (from top to bottom: ECG lead II; MAP 1-2 bipolar electrogram 1.33mV scale; MAP 1 UE 3.45mV scale; CS 1-2 and 5-6 bipolar electrograms); 24-hour clock timeline shown below.

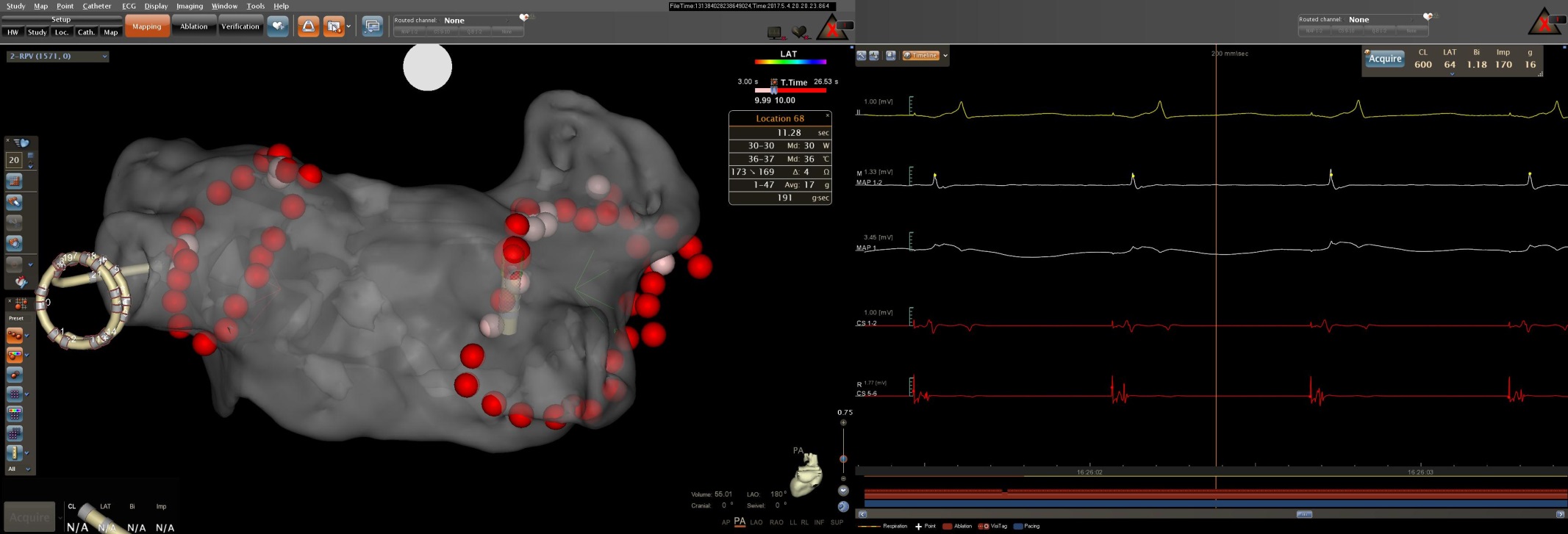

Figure 6B: CARTO^®^3 LA geometry (left panel, transparent PA view) and CARTOREPLAY™ screen (right panel) at the transition to the second annotated RF application during right PV isolation in patient #21 (interval = 17ms indicated by the gap in the red annotation bar at the bottom of the right panel). VISITAG™ Module ablation site annotation is displayed as 2mm radius spheres, with transition to red at 10s RF duration; “Location 68” is highlighted with annotated ablation data shown (box), at a distance of 5.0mm from “Location 67”. CARTOREPLAY™ electrogram data at 200ms sweep speed demonstrates immediate pure R UE morphology of the distal ablation catheter electrogram during CS pacing at 600ms cycle length (from top to bottom: ECG lead II; MAP 1-2 bipolar electrogram 1.33mV scale; MAP 1 UE 3.45mV scale; CS 1-2 and 5-6 bipolar electrograms); 24-hour clock timeline shown below.

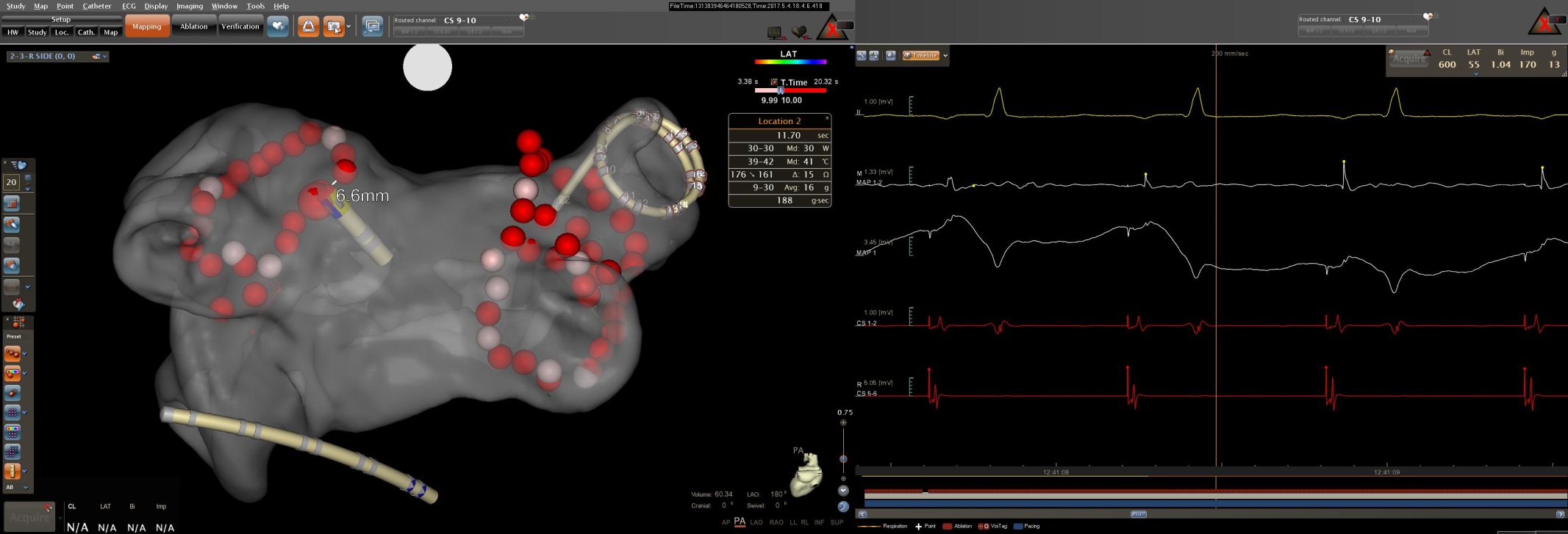

Figure 7A: CARTO^®^3 LA geometry (left panel, transparent PA view) and CARTOREPLAY™ screen (right panel) at the transition to the second annotated RF application during left PV isolation in patient #22 (interval = 17ms indicated by the gap in the red annotation bar at the bottom of the right panel). VISITAG™ Module ablation site annotation is displayed as 2mm radius spheres, with transition to red at 10s RF duration; “Location 2” is highlighted with annotated ablation data shown (box), at a distance of 6.6mm from “Location 1”. CARTOREPLAY™ electrogram data at 200ms sweep speed demonstrates some artefact, but with immediate pure R UE morphology of the distal ablation catheter electrogram during CS pacing at 600ms cycle length still clearly visible (from top to bottom: ECG lead II; MAP 1-2 bipolar electrogram 1.33mV scale; MAP 1 UE 3.45mV scale; CS 1-2 and 5-6 bipolar electrograms); 24-hour clock timeline shown below.

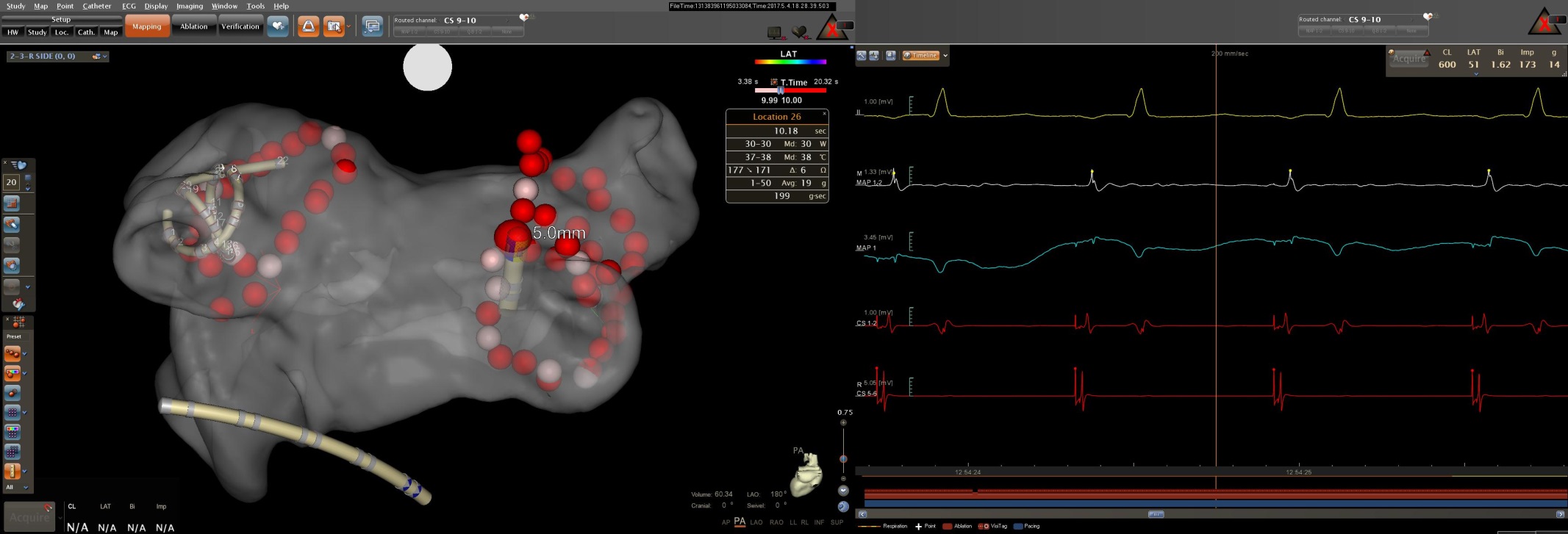

Figure 7B: CARTO^®^3 LA geometry (left panel, transparent PA view) and CARTOREPLAY™ screen (right panel) at the transition to the second annotated RF application during right PV isolation in patient #22 (interval = 16ms indicated by the gap in the red annotation bar at the bottom of the right panel). VISITAG™ Module ablation site annotation is displayed as 2mm radius spheres, with transition to red at 10s RF duration; “Location 26” is highlighted with annotated ablation data shown (box), at a distance of 5.0mm from “Location 25”. CARTOREPLAY™ electrogram data at 200ms sweep speed demonstrates 2 consecutive RS UE morphology distal ablation catheter electrogram complexes during CS pacing at 600ms cycle length, but with the first of 3 consecutive pure R morphology complexes at 1.53s following annotation onset (from top to bottom: ECG lead II; MAP 1-2 bipolar electrogram 1.33mV scale; MAP 1 UE 3.45mV scale; CS 1-2 and 5-6 bipolar electrograms); 24-hour clock timeline shown below.

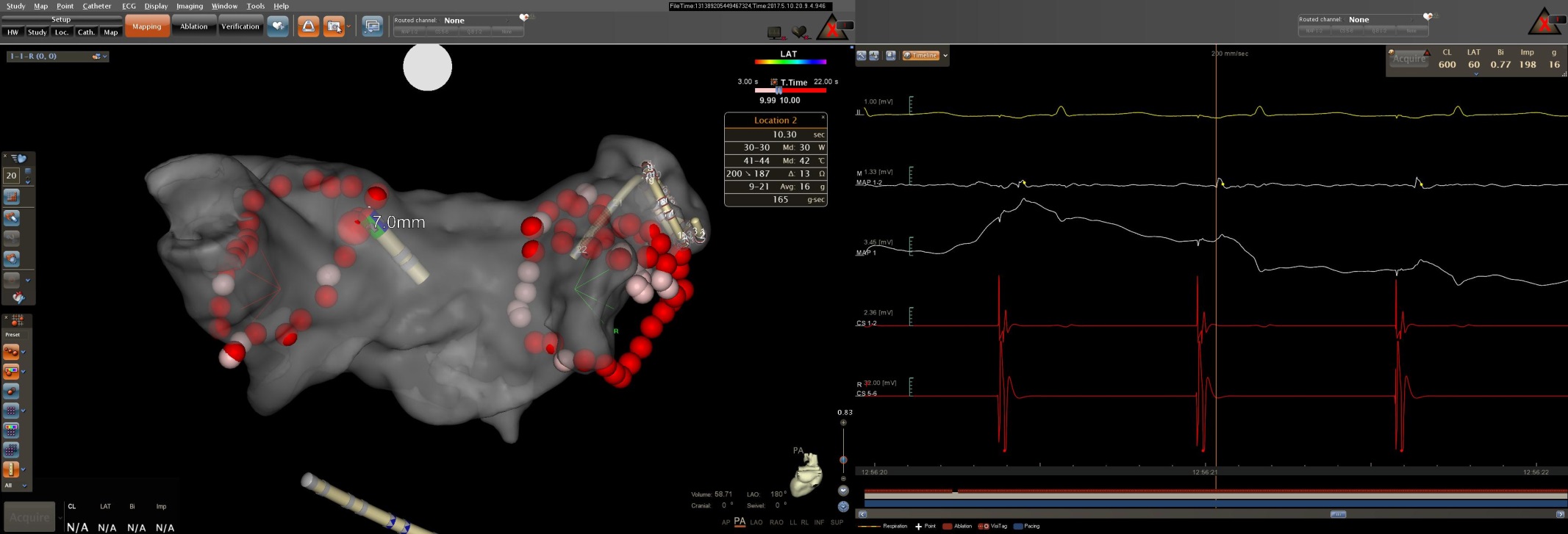

Figure 8A: CARTO^®^3 LA geometry (left panel, transparent PA view) and CARTOREPLAY™ screen (right panel) at the transition to the second annotated RF application during left PV isolation in patient #24 (interval = 17ms indicated by the gap in the red annotation bar at the bottom of the right panel). VISITAG™ Module ablation site annotation is displayed as 2mm radius spheres, with transition to red at 10s RF duration; “Location 2” is highlighted with annotated ablation data shown (box), at a distance of 7.0mm from “Location 1”. CARTOREPLAY™ electrogram data at 200ms sweep speed demonstrates immediate pure R UE morphology of the distal ablation catheter electrogram during CS pacing at 600ms cycle length (from top to bottom: ECG lead II; MAP 1-2 bipolar electrogram 1.33mV scale; MAP 1 UE 3.45mV scale; CS 1-2 and 5-6 bipolar electrograms); 24-hour clock timeline shown below.

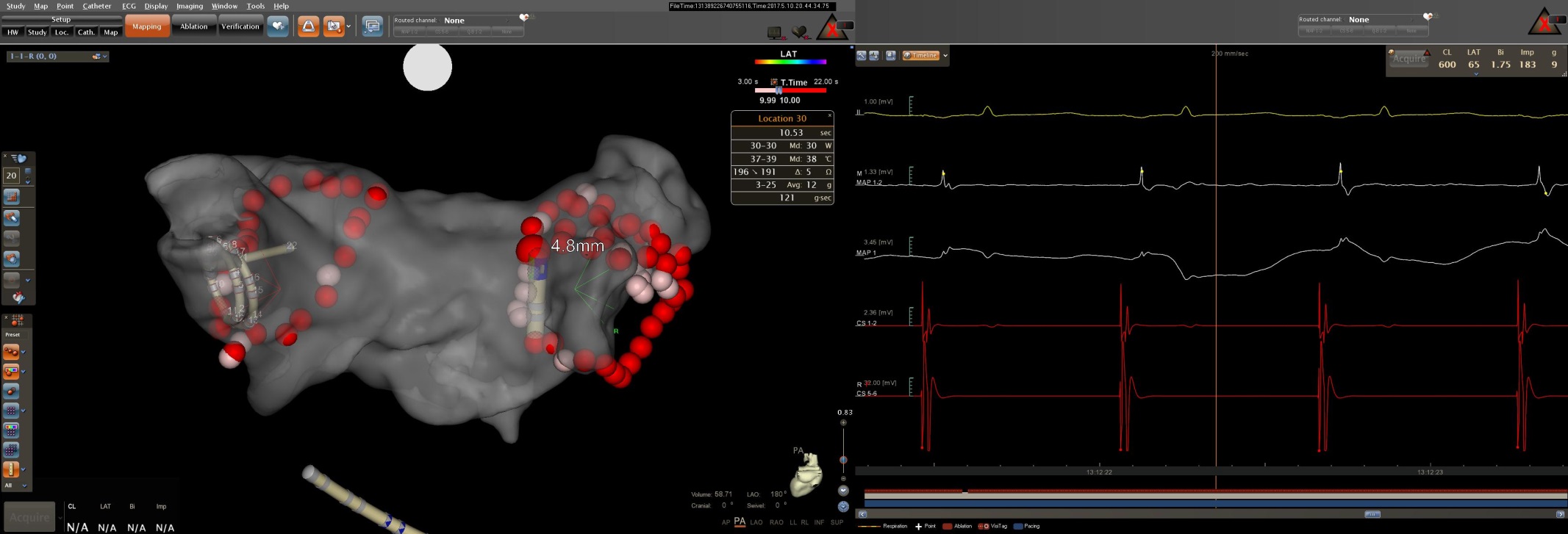

Figure 8B: CARTO^®^3 LA geometry (left panel, transparent PA view) and CARTOREPLAY™ screen (right panel) at the transition to the second annotated RF application during right PV isolation in patient #24 (interval = 16ms indicated by the gap in the red annotation bar at the bottom of the right panel). VISITAG™ Module ablation site annotation is displayed as 2mm radius spheres, with transition to red at 10s RF duration; “Location 30” is highlighted with annotated ablation data shown (box), at a distance of 4.8mm from “Location 29”. CARTOREPLAY™ electrogram data at 200ms sweep speed demonstrates immediate pure R UE morphology of the distal ablation catheter electrogram during CS pacing at 600ms cycle length (from top to bottom: ECG lead II; MAP 1-2 bipolar electrogram 1.33mV scale; MAP 1 UE 3.45mV scale; CS 1-2 and 5-6 bipolar electrograms); 24-hour clock timeline shown below.
